# HGFAC is a ChREBP Regulated Hepatokine that Enhances Glucose and Lipid Homeostasis

**DOI:** 10.1101/2021.07.29.454308

**Authors:** Ashot Sargsyan, Ludivine Doridot, Sarah A. Hannou, Wenxin Tong, Harini Srinivasan, Rachael Ivison, Ruby Monn, Henry H. Kou, Jonathan M. Haldeman, Michelle Arlotto, Phillip J. White, Paul A. Grimsrud, Inna Astapova, Linus Tsai, Mark A. Herman

**Affiliations:** Duke Molecular Physiology Institute, Duke University, Durham, North Carolina, USA; Division of Endocrinology, Beth Israel Deaconess Medical Center, Harvard University, Boston, Massachusetts, USA; Harvard Medical School, Boston, Massachusetts, USA; Division of Endocrinology, Metabolism, and Nutrition, Duke University, Durham, North Carolina, USA; Department of Pharmacology and Cancer Biology, Duke University, Durham, NC, USA; Current affiliation: Université de Paris, Institut Cochin, INSERM, CNRS, F-75014 PARIS, France

## Abstract

Carbohydrate Responsive Element-Binding Protein (ChREBP) is a carbohydrate sensing transcription factor that regulates both adaptive and maladaptive genomic responses in coordination of systemic fuel homeostasis. Genetic variants in the ChREBP locus associate with diverse metabolic traits in humans, including circulating lipids. To identify novel ChREBP-regulated hepatokines that contribute to its systemic metabolic effects, we integrated ChREBP ChIP-seq analysis in mouse liver with human genetic and genomic data for lipid traits and identified Hepatocyte Growth Factor Activator (HGFAC) as a promising ChREBP-regulated candidate in mice and humans. HGFAC is a protease that activates the pleiotropic hormone Hepatocyte Growth Factor (HGF). We demonstrate that HGFAC KO mice have phenotypes concordant with putative loss-of-function variants in human HGFAC. Moreover, in gain- and loss-of-function genetic mouse models, we demonstrate that HGFAC enhances lipid and glucose homeostasis, in part, through actions to activate hepatic PPARγ activity. Together, our studies show that ChREBP mediates an adaptive response to overnutrition via activation of an HGFAC-HGF-PPARγ signaling axis in the liver to preserve glucose and lipid homeostasis.

## Introduction

Carbohydrate Responsive Element Binding Protein (ChREBP, also known as Mlxipl) is a transcription factor expressed in key metabolic tissues including liver, adipose tissue, kidney, small intestine, and pancreatic islets (1, 2). It is activated by sugar metabolites, and in the liver and small intestine, its activity is robustly increased by consumption of sugars containing fructose (3, 4). Upon activation, ChREBP stimulates expression of genomic programs that contribute to both adaptive and maladaptive metabolic responses (1). Hepatic ChREBP activity is increased in human obesity and diabetes as indicated by increased expression of the potent ChREBP-beta isoform (5, 6). Knockdown or knockout of hepatic ChREBP protects against diet-induced and genetic forms of obesity and accompanying metabolic disease (3, 7, 8).

ChREBP plays a significant role in human metabolic physiology as common genetic variants in the ChREBP locus associate with pleotropic anthropomorphic and metabolic traits at genome-wide significance including circulating lipids and cholesterol, BMI, waist-hip ratio, height, diverse hematological parameters, serum urate, liver enzymes, and blood pressure (9). However, the complement of ChREBP transcriptional targets that participate in regulating these diverse traits is incompletely understood. To date, thousands of genes have been proposed as putative ChREBP targets via Chromatin Immunoprecipitation-sequencing (ChIP-seq) assays and global gene expression analysis, a minority of which are known to be involved in metabolic processes (10, 11). For example, it is well established that ChREBP regulates glycolysis and fructolysis, hepatic and adipose lipogenesis, and hepatic glucose production via regulation of key enzymes involved in these metabolic pathways (4, 12–14). At the same time, a large number of putative ChREBP transcriptional targets have either no known or poorly defined function and uncertain impact on metabolism.

Here we performed ChIP-seq for ChREBP in mouse liver and integrated this with human genetic data to identify novel ChREBP-dependent hepatokines that might participate in the regulation of systemic metabolism. Through this screen we identified Hepatocyte Growth Factor Activator (HGFAC) as a promising candidate. HGFAC is a liver-secreted, circulating protease that activates Hepatocyte Growth Factor (HGF) which has pleiotropic biological activities including regulation of morphogenesis, cell migration, transition between cell states, and proliferation in epithelial and other cell types throughout the body (15–17). Here, we demonstrate that HGFAC is indeed nutritionally regulated in a ChREBP-dependent manner and participates in an adaptive response to preserve carbohydrate and lipid homeostasis.

## Results

### HGFAC is a ChREBP genomic target that associates with metabolic traits in humans

To identify ChREBP transcriptional targets that participate in the regulation of ChREBP associated metabolic programs and phenotypes, we performed ChIP-seq analysis for ChREBP in livers of two strains of male mice gavaged with either water or fructose. We identified 4,860 distinct genomic sites enriched for ChREBP binding (**Supplementary Table 1**) which include well-defined loci in canonical ChREBP targets involved in glycolysis, glucose production, fructolysis, and lipogenesis such as liver pyruvate kinase (PKLR), glucose-6-phosphatase (G6PC), fatty acid synthase (FASN), and ketohexokinase (KHK), respectively (**Figure 1A**). Although fructose gavage can acutely induce ChREBP-dependent changes in gene expression, ChREBP ChIP-seq peaks were readily detectable in fasted mice, and fructose gavage did not enhance ChREBP ChIP-seq peak height even at a liberal false discovery rate of 0.20. This indicates that increased chromatin occupancy is not essential for fructose to induce ChREBP-dependent gene transcription. Most ChREBP ChIP peaks occurred within 10 kb of transcriptional start sites (**Figure 1B**). Consistent with ChREBP’s known functions, Genomic Region Enrichment Analysis (GREAT) of putative ChREBP binding sites demonstrated enrichment for numerous metabolic processes including carbohydrate and lipid metabolism (**Figure 1C**) (18).

**Figure 1.**
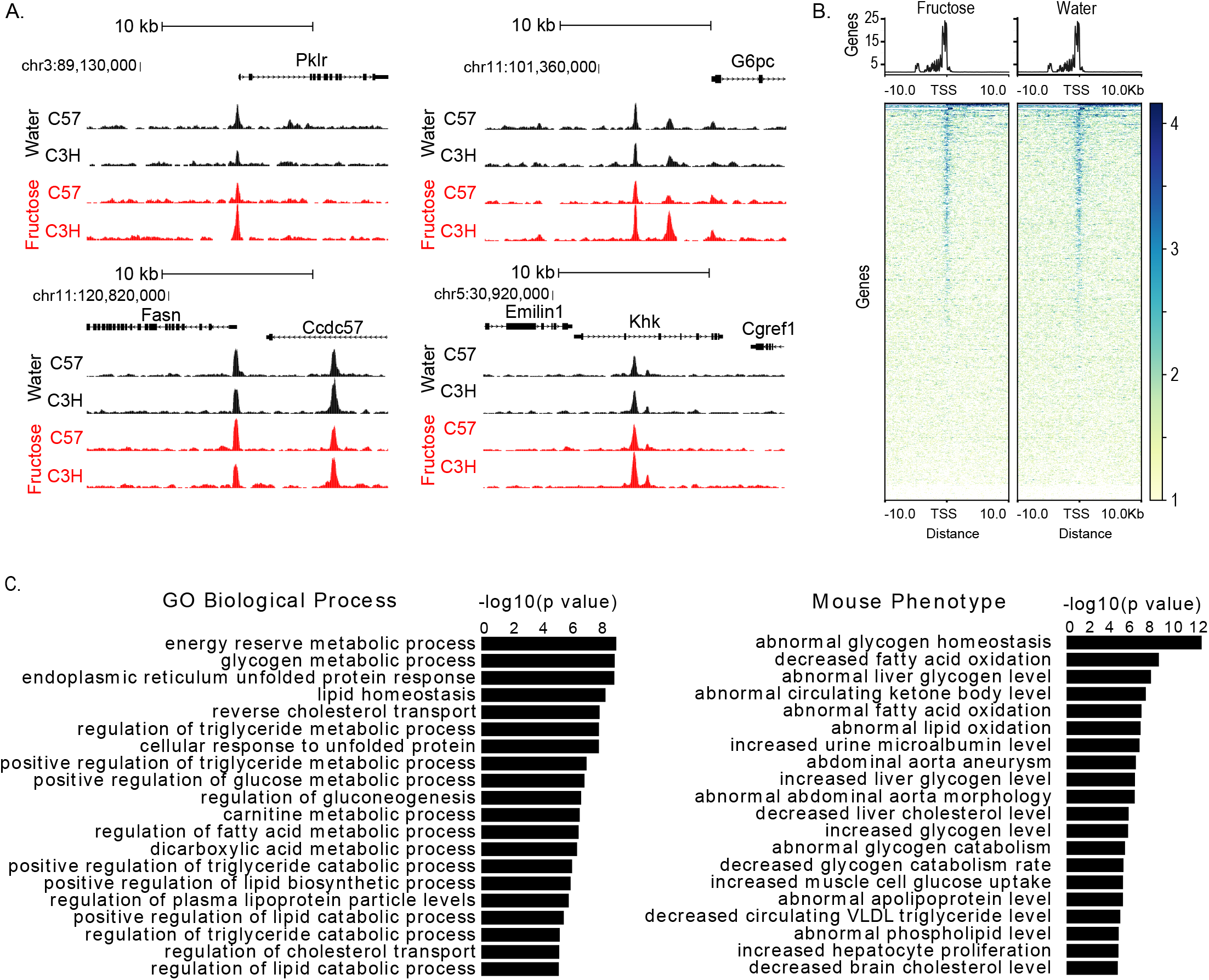
ChREBP is bound to liver genomic targets following water and fructose gavage after a 5 hour fast. A) ChREBP ChIP-seq signal tracks in liver of male C57 and C3H mice after a 5 hour fast and 90 minutes after water versus fructose gavage (4 g/kg body weight) in selected ChREBP transcriptional targets including Pklr, Gp6c, Fasn, and Khk. B) Heatmaps showing hepatic ChREBP peaks after fructose versus water gavage. The amplitude of each peak center is represented by the z score and shown in blue. C) “GO Biological Process” and “Mouse Phenotype” pathway analysis for ChREBP peaks.

Variants in the ChREBP locus are strongly associated with hypertriglyceridemia in human populations (19, 20). However, the complement of ChREBP transcriptional targets that mediate its effects on circulating lipids is uncertain. We sought to determine whether genomic loci containing human homologues of mouse ChREBP target genes are enriched for variants that associate with hypertriglyceridemia in human populations. Via Meta-Analysis of Gene-set ENrichmenT of variant Associations (MAGENTA), we confirmed that loci in proximity to human homologues of mouse genes that are within 20 kb of ChREBP binding sites are enriched for SNPs that associate with hypertriglyceridemia in humans (Adjusted P-val = 0.003). 87 loci/genes contributed to this enrichment with an FDR of 0.05 (**Table 1 and Supplementary Table 2**) (21). This list includes known ChREBP transcriptional targets such as *GCKR, TM6SF2, KHK*, and *ChREBP* (*MLXIPL*) itself, all previously implicated in regulating carbohydrate and triglyceride metabolism. Of these 87 loci, seven encoded putative secretory proteins including several lipoproteins (APOC2, APOE, and APOA5), VEGFA which is most well-known for its role in angiogenesis, but also implicated in metabolic control, and HGFAC (**Supplementary Table 2)** (22). To our knowledge, HGFAC has not been identified as a ChREBP transcriptional target nor studied extensively in the context of systemic fuel metabolism.

**Table 1.** Top 25 gene candidates in proximity to ChREBP ChIP-seq binding sites and that contribute to enrichment for association with hypertriglyceridemia as assessed by MAGENTA.

| Gene Symbol | Entrez ID | Gene<br>-log<sub>10</sub>(p-value) | Best SNP<br>rsID | Best SNP<br>-log<sub>10</sub>(pval) |
| --- | --- | --- | --- | --- |
| PREB | 10113 | 11 | rs1659689 | 33.9 |
| APOC2 | 344 | 10.8 | rs439401 | 65.8 |
| GCKR | 2646 | 10.5 | rs1260326 | 238.6 |
| REEP3 | 221035 | 10.4 | rs7897379 | 16.9 |
| APOE | 348 | 10.4 | rs439401 | 65.8 |
| KHK | 3795 | 10.1 | rs7588926 | 14.9 |
| JMJD1C | 221037 | 10.1 | rs10761762 | 17 |
| LPAR2 | 9170 | 10 | rs10500212 | 59.7 |
| SIK3 | 23387 | 9.9 | rs6589574 | 112.5 |
| APOA5 | 116519 | 9.9 | rs10790162 | 249 |
| MLXIPL | 51085 | 9.9 | rs11974409 | 99.9 |
| TM6SF2 | 53345 | 9.8 | rs10401969 | 69 |
| SLC5A6 | 8884 | 9.6 | rs1659685 | 39.1 |
| NDUFA13 | 51079 | 9.5 | rs3794991 | 62 |
| VPS37D | 155382 | 9.5 | rs7777102 | 89.4 |
| VEGFA | 7422 | 9.4 | rs998584 | 14.5 |
| GRB14 | 2888 | 8.8 | rs10195252 | 14.2 |
| MSL2 | 55167 | 7.5 | rs645040 | 11.7 |
| HGFAC | 3083 | 7.3 | rs6831256 | 11.8 |
| MAFF | 23764 | 6.5 | rs3761445 | 11.1 |
| PLA2G6 | 8398 | 6.4 | rs3761445 | 11.1 |
| STX1A | 6804 | 6.2 | rs1128349 | 11 |
| LRP1 | 4035 | 5 | rs11172134 | 8.8 |
| BCL3 | 602 | 4.5 | rs4803750 | 8 |
| CBLC | 23624 | 4.5 | rs4803750 | 8 |

HGFAC is a serine protease expressed predominately in hepatocytes and secreted as a zymogen into circulation where it is found in a single chain pro-HGFAC form (23, 24). In-vitro studies have identified thrombin and kallikrein-related peptidases KLK-4 and KLK-5 to be potent activators of pro-HGFAC (25, 26). Once activated, HGFAC cleaves and activates Hepatocyte Growth Factor (HGF) which can then bind and activate the c-Met receptor tyrosine kinase (MET) (23). HGF and c-MET have pleiotropic biological activities as mitogens and motogens in organogenesis, tissue repair, and cell migration, and also function as anti-inflammatory, apoptotic, and cytoprotective signals depending on the context (15). Variants in *c-MET* also associate with circulating triglycerides at genome wide significance in humans consistent with a potential role for HGFAC in regulating triglyceride levels through activation of HGF (27). Moreover, increased levels of circulating HGF in people associate with features of cardiometabolic disease including obesity, risk for type 2 diabetes, and risk for cardiovascular disease (28–31). Circulating HGF levels are influenced by variants in the HGFAC locus (32). A missense variant in HGFAC, rs3748034, that associates with increased circulating HGF also associates with increased circulating triglycerides in GWAS aggregate data at genome wide significance (beta = 0.0302, p < 5e-28) as well as other cardiometabolic risk factors and pleiotropic biological traits (33, 34). The Ala218Ser mutation encoded by rs3748034 is predicated to be “possibly damaging” by PolyPhen-2 (35). Furthermore, another missense variant in HGFAC, rs16844401, that associates with increased circulating triglycerides also associates with increased coronary artery disease risk (36). These associations motivated further investigation to determine whether ChREBP regulates HGFAC expression and whether this interacts with nutritional status to regulate systemic fuel metabolism and cardiometabolic risk factors.

### Nutritional regulation of HGFAC is ChREBP-dependent

ChREBP activity in the liver is responsive to diets high in fructose. To examine the role of hepatic ChREBP in the regulation of HGFAC in rodents, we measured hepatic *Hgfac* mRNA and HFGAC protein in the liver and plasma of mice with liver specific deletion of ChREBP after 8 weeks on high fructose (HFrD) or control diet. High fructose feeding increased hepatic *Hgfac* mRNA expression 1.7-fold (p<.0001) in control mice, and this induction was abrogated in liver-specific ChREBP KO mice (LiChKO) (**Figure 2A**). Fructose-induced increases in hepatic *Hgfac* mRNA expression were accompanied by 4- and 2-fold increases in hepatic and circulating pro-HGFAC protein levels (**Figure 2B**). Basal liver and circulating HGFAC protein levels tended to be decreased in chow fed LiChKO mice and were not induced with fructose feeding. Circulating HGFAC also increased in mice fed a high fat/high-sucrose (HF/HS) diet and in genetically obese Zucker fatty rats on chow diet (**Supplementary Figure 1**), where hepatic ChREBP activity is also robustly increased independently of an obesogenic diet (37). Altogether, these data show that hepatic ChREBP mediates diet and obesity induced increases in circulating HGFAC.

**Figure 2.**
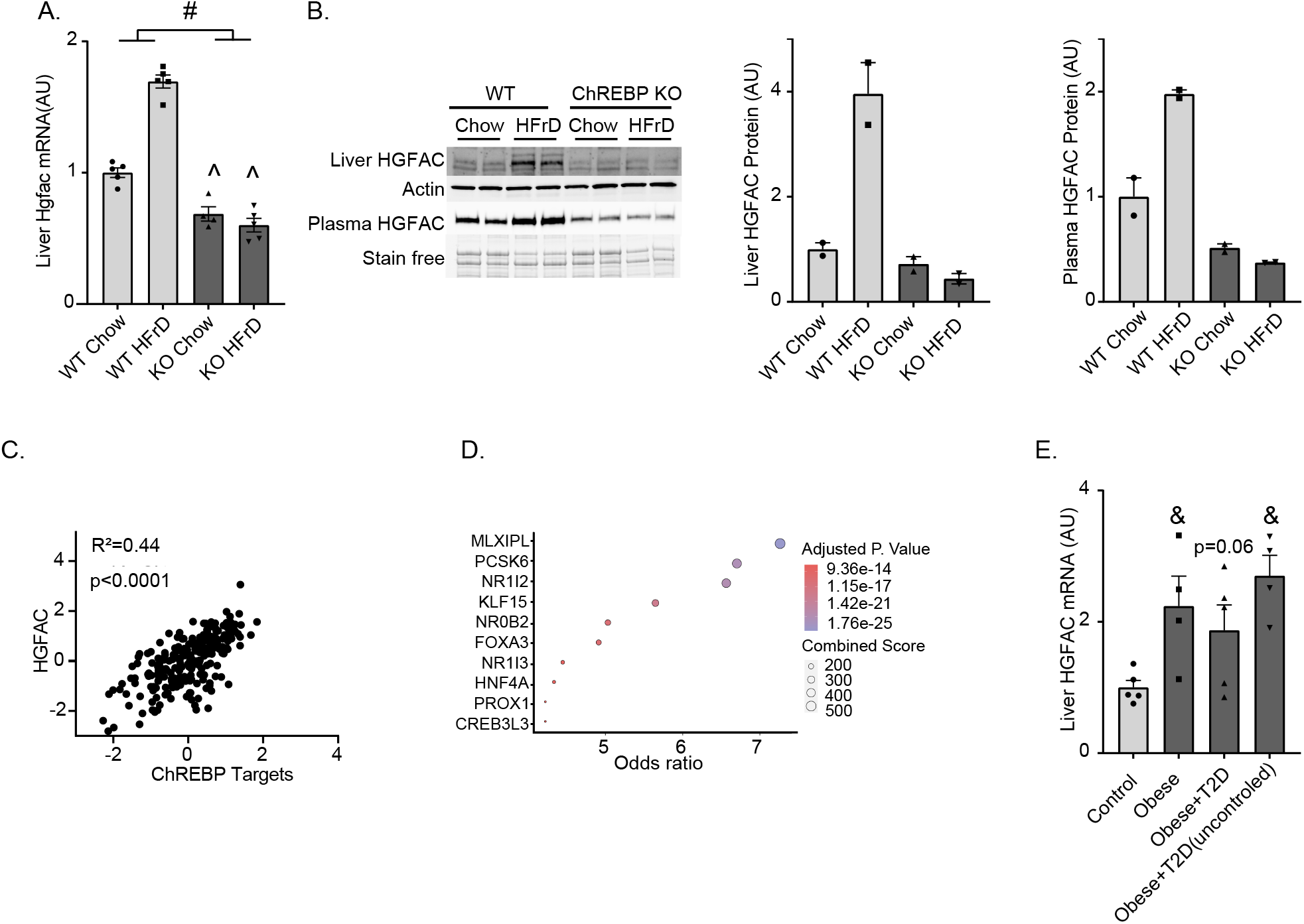
ChREBP Links Nutritional Status to Circulating HGFAC. A) Liver mRNA expression and B) liver and circulating levels of HGFAC protein in WT (wild-type, littermate control) and KO (liver specific ChREBP KO) mice after 8 weeks on chow versus high fructose diet (HFrD) with their quantification by densitometry. C) Correlation between *HGFAC* mRNA expression and a composite vector comprised of canonical ChREBP transcriptional targets in human livers from the GTEx project (Pearson correlation R^2^=0.44, p<0.0001, n=226). D) Factors ranked by odds ratio for enrichment of the 300 genes most highly co-expressed with the factor in the ARCHS4 project that are also present in the top 5% of genes that correlate with HGFAC expression in the GTEx project. Combined score = log(p)*z, where p is calculated by Fisher’s exact test and z is the z-score calculated by assessing the deviation from the expected rank. The size of the circle corresponds to the enrichment score and the color corresponds to the adjusted p-value. E) Expression of *HGFAC* mRNA in livers of healthy controls, obese non-diabetic subjects, obese subjects with well controlled diabetes, and obese subjects with poorly controlled diabetes, n=4-5 per group. Data represent means ± SEM. Statistics were assessed by 2-way ANOVA with Sidak’s multiple comparisons between individual groups, # p<0.05, for main effects, ^ p<0.05 for comparison across genotypes within diets; or one-way ANOVA with Holm-Sidak’s multiple comparisons test between control and other groups, & p<0.05.

Next, we sought to determine whether ChREBP-mediated regulation of HGFAC might be conserved in humans. To that end, we analyzed hepatic mRNA expression levels of *HGFAC* and other ChREBP transcriptional targets in the Genotype-Tissue Expression (GTEx) Biobank (38). Expression of the potent *ChREBP-*β isoform is an excellent surrogate marker of tissue ChREBP activity (14). However, it is expressed at low levels which are typically below the sequencing depth of most RNA-seq experiments. Consistent with this, GTEX RNA-seq data does not distinguish between *ChREBP*-β and -α isoforms. Due to the lack of *ChREBP-*β specific expression data, we used a composite expression vector comprised of 5 well-validated ChREBP target genes (*FASN, PKLR, KHK, ALDOB, and SLC2A2*) and found that this composite vector strongly correlates with the expression of *HGFAC* (Pearson correlation r^2^=0.44, p <0.0001) **(Figure 2C)**. Transcription factor enrichment analysis of the 5% of hepatic genes that best correlated with hepatic *HGFAC* expression in the GTEx Biobank showed strong enrichment for genes co-expressed with ChREBP (Adjusted P-val = 1.75E-25) indicating conservation of the ChREBP-mediated regulation of HGFAC in humans **(Figure 2D)** (39). Additionally, hepatic *HGFAC* mRNA expression is upregulated in patients with obesity and uncontrolled diabetes **(Figure 2E)**, conditions that are associated with increased hepatic ChREBP activity (5, 40). Collectively, these data support the hypothesis that hepatic HGFAC expression and circulating levels of HGFAC are regulated by ChREBP activity both in rodents and in humans, and hepatic *HGFAC* expression is increased in obesity and diabetes.

### Murine Hgfac KO recapitulates the phenotype of putative human LOF HGFAC variants

To study the roles of HGFAC in systemic metabolism, we generated whole body *Hgfac* KO mice that lack a portion of exon 1 and all of exon 2 **(Figure 3A)**. The deletion was confirmed by genomic PCR, by the absence of detectable circulating HGFAC protein, and by quantification of hepatic *Hgfac* mRNA **(Figure 3B-D)**. *Hgfac* KO mice were born at normal Mendelian ratios and did not appear to have any gross abnormalities when compared to their littermate controls. Activated HGFAC activates HGF and c-MET signaling. However, there is redundancy in this system and other proteases including Hepsin (HPN) and coagulation factors XIa and XIIa are also capable of activating HGF (41, 42). We sought to determine whether the ability to activate endogenous HGF is impaired in *Hgfac* KO mice. c-MET signaling was assessed in HepG2 cells incubated with thrombin treated sera obtained from WT and *Hgfac* KO mice. Thrombin is one of the proteases that is capable of activating HGFAC *in-vitro* (26). Serum from control mice increased c-MET phosphorylation 1.9-fold when compared to DMEM alone, while this induction was attenuated with serum from KO mice **(Figure 3E)**. These results demonstrate that sera from *Hgfac* KO mice has reduced capacity to activate HGF and c-MET signaling.

**Figure 3.**
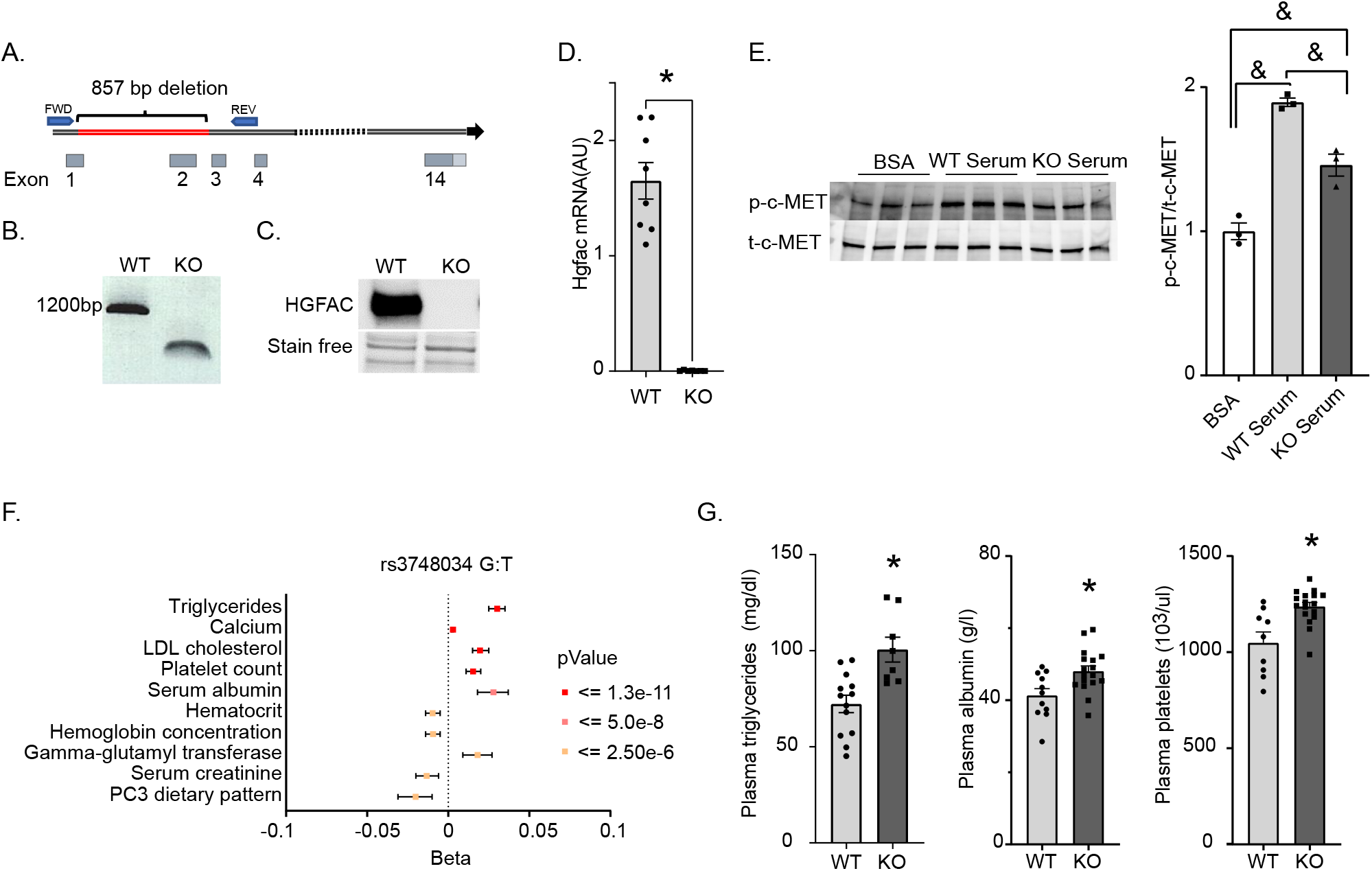
The phenotype in HGFAC KO mice recapitulates the phenotype of putative loss of function variant in human HGFAC. A) Schematic depiction of *Hgfac* gene and the deleted region in red; FWD and REV indicate the positions of forward and reverse primers, respectively, used in genomic PCR shown in (B) confirming the deletion of an 857 bp region in the HGFAC gene. C) Representative immunoblot of circulating HGFAC in WT (wild-type, littermate control) and KO (HGFAC KO) plasma. D) Hepatic *Hgfac* mRNA levels measured by qPCR in WT and HGFAC KO mice, n=7-9/group. E) Immunoblot and quantification of phospho-c-MET in HepG2 cells treated with activated sera of WT and HGFAC KO mice, n=3 per condition. F) Forest plot of phenotypes associated with the rs3748034 putative loss of function coding variant in human HGFAC. G) Quantification of plasma triglyceride levels in ad libitum fed male WT and HGFAC KO mice, n=8-13 / group, plasma albumin concentrations in male WT and HGFAC KO mice, n=11-17 / group, and plasma platelet levels in male WT and HGFAC KO mice, n=9-17 / group. Data represent means ± SEM, Statistics were assessed by two-tailed unpaired t-test, * p<0.05; or one-way ANOVA with Holm-Sidak’s multiple comparisons test between groups, & p<0.05.

A putative loss of function variant in *HGFAC* (rs3748034) strongly associates with increased circulating triglycerides, albumin, and platelets among other traits (**Figure 3F**) (33). We determined whether *Hgfac* KO mice have similar phenotypes. *Hgfac* KO mice had a 28% increase in circulating triglycerides (100 +/− 6.5 mg/dl vs 72+/− 4.5 mg/dl, p<0.001), a 15% increase in circulating albumin (4.8 +/− 0.19 vs 4.1 +/− 0.15 g/dl, p<0.01), and a 15% increase in circulating platelets (1237 +/− 22 cells*10^3^/ul vs 1048 +/− 57 cells*10^3^/ul, p<0.05) (**Figure 3G**). No hematological parameter other than platelet count was altered **(Supplementary Figure 2)**. Collectively, these data indicate that murine *Hgfac* KO recapitulates phenotypes in putative loss-of-function human *HGFAC* variants.

### HGFAC KO Mice develop impaired glucose homeostasis

To examine the potential role of HGFAC in systemic metabolism, we challenged 8-week-old *Hgfac* KO mice and their littermate controls with high-fat/high-sucrose (HFHS) diet for 18 weeks. We did not observe any differences in body weight or fat mass during the study **(Figure 4A and B)**. However, a modest reduction in lean body mass was observed in *Hgfac* KO mice **(Figure 4C)**. To assess glucose homeostasis, we performed glucose and glycerol tolerance tests in *Hgfac* KO mice and controls at time points throughout the study. Glycerol is a preferred gluconeogenic substrate and glycerol tolerance tests reflect hepatic glucose production capacity (43). After 4 weeks on HFHS diet, *Hgfac* KO mice are glycerol intolerant with a 1.4-fold increase in glycemic excursion (p<0.05) **(Figure 4D)**. At this age, there was no difference between KO mice and controls with respect to glycemic excursion during a glucose tolerance test **(Figure 4E)**, suggesting that young *Hgfac* KO animals may have a selective impairment in hepatic insulin sensitivity. However, after 13 weeks of HFHS diet, *Hgfac* KO mice developed glucose intolerance with a 1.6-fold increase in incremental AUC (p<0.005) as well as insulin resistance with a 30% decrease in area above the curve (p<0.05), as measured by IP glucose and insulin tolerance tests, respectively (**Figure 4F and G)**.

**Figure 4.**
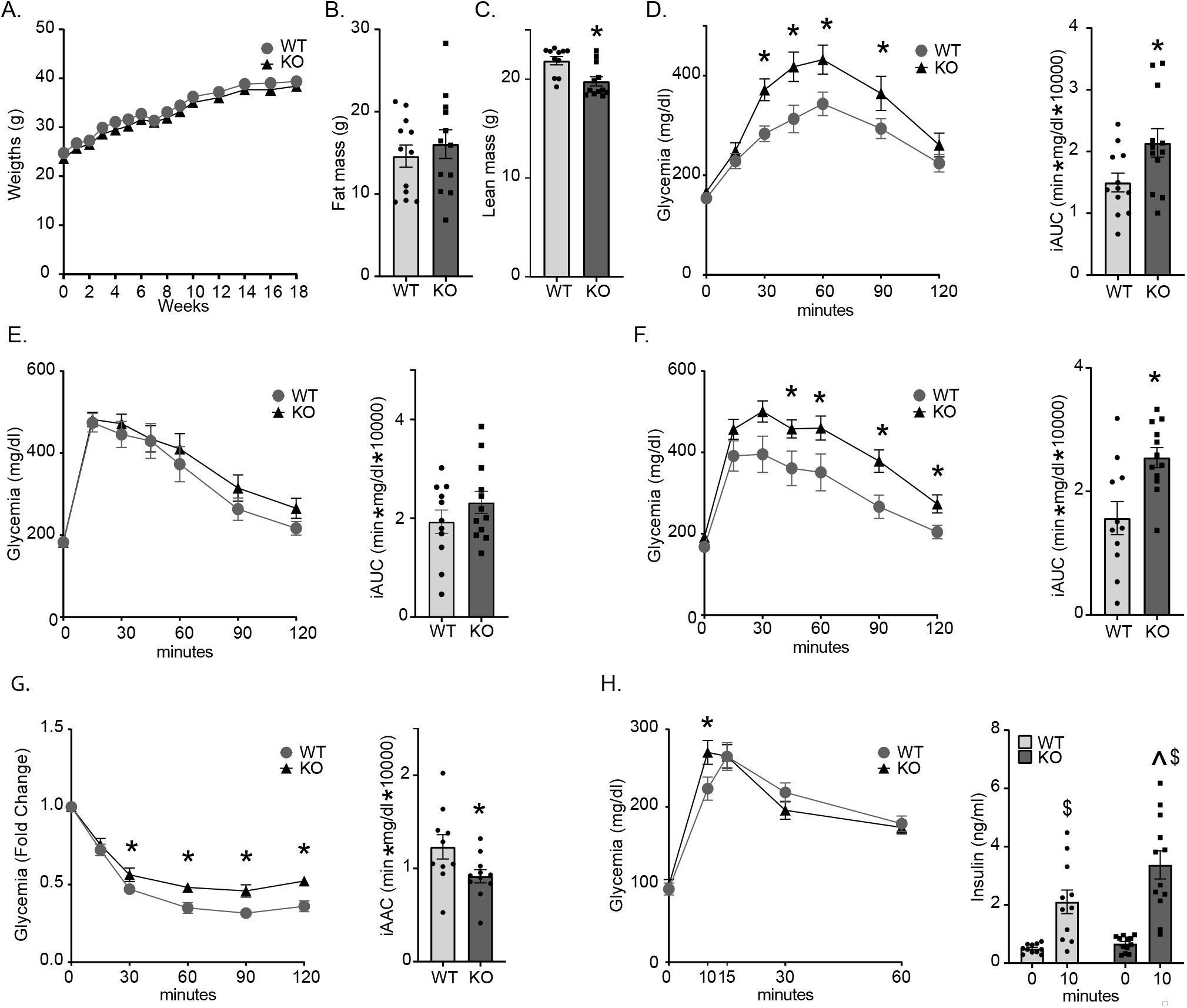
HGFAC KO mice have impaired carbohydrate metabolism on HF/HS diet. A) Body weight of male WT and HGFAC KO mice during 18 weeks of HF/HS feeding (n=11-12/group unless otherwise specified) B) Fat and C) lean mass by NMR at 18 weeks. Glucose homeostasis was assessed at intervals throughout the study including D) IP glycerol tolerance test at 4 weeks, E) IP glucose tolerance at 5 weeks, F) IP glucose tolerance test at 13 weeks, G) IP insulin tolerance test at 14 weeks, and H) a mixed meal tolerance test to assess insulin secretion was performed at 16 weeks. Tail vein insulin levels were measured at 0 and 10 minutes. Data represent means ± SEM, Statistics were assessed by two-tailed unpaired t-test, *p<0.05; or two-way ANOVA with Sidak’s multiple comparisons between individual groups, ^ p<0.05 for comparison across genotypes within time points, $ p<0.05 for comparison across time points within genotypes.

HGF has been proposed to regulate pancreatic beta-cell development and insulin secretory capacity (44). To test insulin secretory capacity in *Hgfac* KO mice, we performed an oral mixed meal tolerance test which triggers more robust and sustained insulin secretion compared to IP glucose administration. Basal insulin and glucose levels were no different between *Hgfac* KO mice and controls **(Figure 4H)**. At 10 min, insulin levels were 1.6-fold higher in *Hgfac* KO mice compared to controls (3.37+/− 0.48 ng/ml *Hgfac* KO vs 2.1+/− 0.4 ng/ml controls, p<0.05) with only a modest increase in glycemia at this time point. Altogether, these data indicate that *Hgfac* KO mice subjected to HFHS diet develop early hepatic insulin resistance followed by systemic insulin resistance with intact insulin secretory capacity.

We also examined whether HFHS diet might exacerbate the increase in circulating triglyceride levels observed in chow-fed *Hgfac* KO mice. In contrast with the data in chow diet, we did not observe differences in circulating triglycerides after 7 weeks of HFHS diet. Similarly, triglyceride levels were not different between *Hgfac* KO and control after IP administration of poloxamer 407 which inhibits lipoprotein lipase and peripheral triglyceride clearance (**Supplementary Figure 3**) indicating that VLDL production is similar between genotypes in this dietary context (45).

### Hgfac KO Downregulates Hepatic PPARγ Expression

To define mechanisms that might contribute to altered triglyceride and carbohydrate metabolism in *Hgfac* KO mice, we performed RNA-seq analysis on liver from chow and HFHS-fed *Hgfac* KO mice and littermate controls after 4 weeks on diet. *Hgfac* was the most significantly downregulated mRNA on both diets, confirming successful KO **(Figure 5A)**. By pathway enrichment analysis **(Figure 5B)**, genes involved in cell cycling were the most downregulated set in chow-fed *Hgfac* KO mice. This is consistent with HGF’s known effects to stimulate hepatocyte proliferation (46). Pathway analysis also suggested changes in lipid metabolism with reduced “PPAR signaling pathway” and “Fatty acid degradation” in KO mice on both diets. Upregulation of genes involved in ribosomal function were observed in the *Hgfac* KO mice potentially consistent with reduced cell cycling and enhanced differentiated function as a result of reduced HGF signaling. Gene sets associated with complement and coagulation pathways were also upregulated in *Hgfac* KO mice. Upregulation of complement and coagulation pathways is notable as putative loss of function variants in the *HGFAC* locus also associate with increased circulating fibrinogen levels (47).

**Figure 5.**
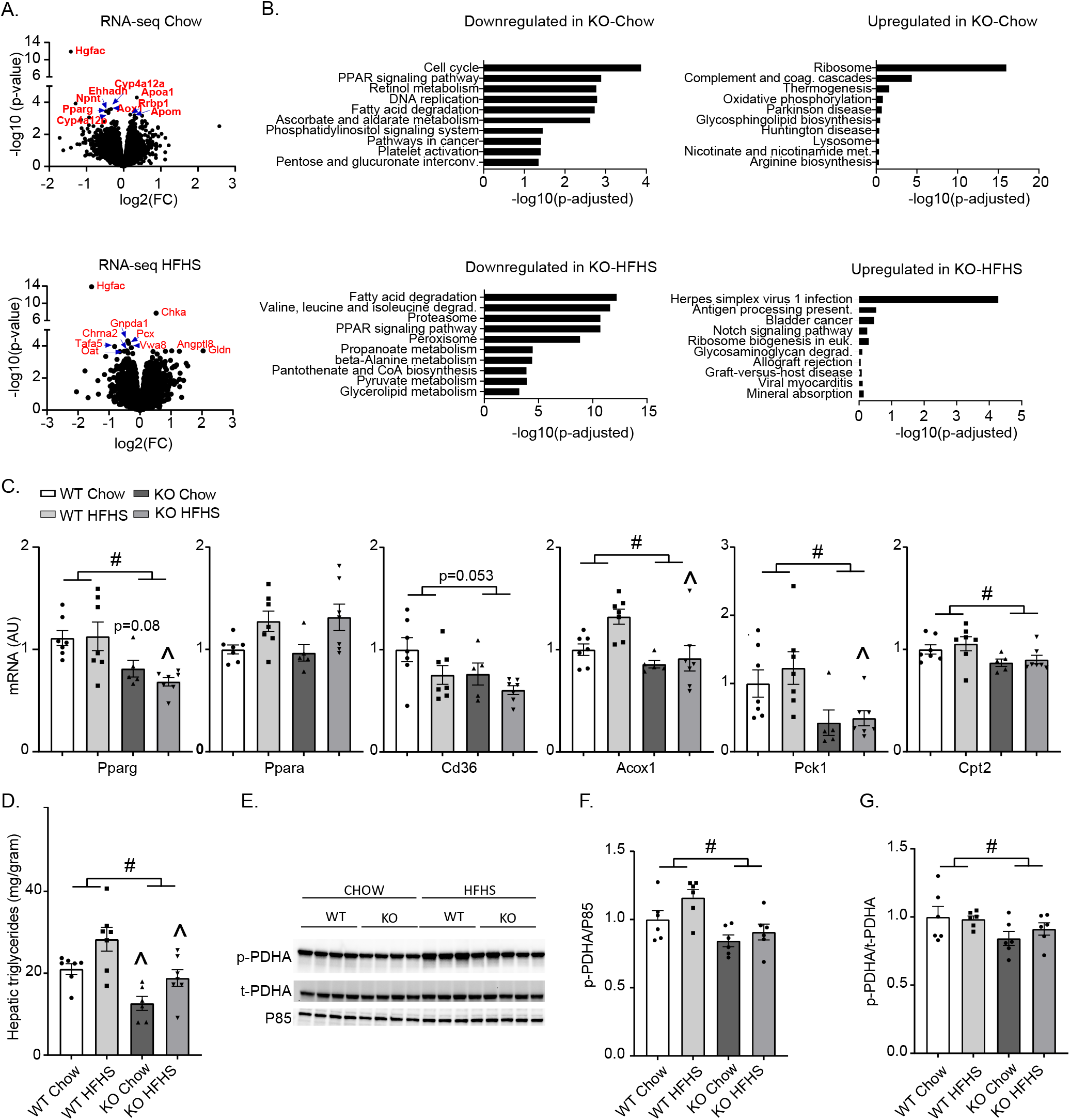
Hepatic Pparg is down-regulated in HGFAC KO mice. A) Volcano plot depicting differentially expressed genes from livers of chow and HF/HS fed HGFAC KO mice versus controls. Named genes in red represent top 10 most differentially expressed genes ranked by p-value. B) Pathway analysis including the top 10 most downregulated and upregulated gene sets, respectively in chow and HF/HS fed HGFAC KO livers compared to controls. C) Hepatic mRNA levels of Pparg, Ppara, Cd36, Acox1, Pck1 and Cpt2 after 4 weeks of Chow or HF/HS diet n=5-7/group. D) Hepatic triglyceride levels in WT and HGFAC KO mice on chow and HF/HS diet after overnight fasting followed by 4-hour ad libitum refeeding (n=6-7/group). E) Immunoblot analysis and quantification of hepatic phospho-S293 PDHA, total PDHA and p85 loading control in chow or HF/HS fed HGFAC KO and controls with quantification normalized to (F) P85 or to (G) total PDHA (n=6/group). Data represent means ± SEM, Statistics were assessed by two-way ANOVA with Sidak’s multiple comparisons between individual groups, ^#^ p<0.05, for genotype main effects, ^ p<0.05 for comparison across genotypes within diets.

Consistent with the pathway analysis, *Pparg* was in the top 10 most differentially expressed genes comparing chow-fed HGFAC KO mice and controls **(Supplementary Table 3).** To confirm this, we quantified hepatic mRNA gene expression by qPCR which revealed that *Pparg* but not *Ppara* is downregulated in livers of chow and HFHS-fed *Hgfac* KO mice compared to controls (**Figure 5C**). Furthermore, PPARγ target genes were also downregulated. These results were replicated in a second cohort (**Supplementary Figure 4**). Hepatic PPARγ is reported to enhance liver fat accretion yet preserve hepatic and systemic insulin sensitivity (48, 49). HFHS-feeding increased the levels of hepatic triglycerides by 49% and 34% in *Hgfac* KO mice and controls, respectively (**Figure 5D**). However, hepatic triglyceride levels were reduced by 40% and 32% in *Hgfac* KO mice compared to controls on chow and HFHS-diets, respectively. Recently, Shannon *et al*. reported that pioglitazone, a PPARγ agonist, increases phosphorylation (S293) of catalytic subunit of hepatic pyruvate dehydrogenase (PDHA), which inhibits PDHA activity and diminishes hepatic glucose output but increases the level of hepatic triglycerides (50). Consistent with this mechanism, we observed reduced S293-PDHA phosphorylation in HGFAC KO animals on chow and HFHS-diets indicative of increased PDHA activity (**Figure 5E-G**). Altogether, phenotypes in *Hgfac* KO mice are consistent with liver-specific deletion of PPARγ which results in reduced hepatic steatosis and impaired hepatic glucose homeostasis eventually leading to the development of peripheral insulin resistance (48, 49). This may be in part mediated by the effects of PPARγ on hepatic PDHA activity.

### HGFAC overexpression enhances glucose homeostasis

As HGFAC deficiency decreased expression of hepatic *Pparg* and its targets, we examined whether HGFAC overexpression has reciprocal molecular and metabolic effects. Adenoviral (ADV) mediated overexpression of HGFAC resulted in a robust increase of circulating HGFAC over a period of two weeks compared to ADV-GFP controls **(Figure 6A)**. This was associated with markedly improved glucose tolerance with a 30% reduction in incremental AUC (p<0.005) **(Figure 6B)** and a 50% reduction in glycemic excursion during a glycerol tolerance test performed in second cohort (p<0.0005) **(Supplementary Figure 5A)**. Additionally, fasting glucose levels were slightly but significantly lower in ADV-HGFAC mice in fed and fasted conditions (**Supplementary Figure 5B)**. Analysis of hepatic gene expression revealed that HGFAC overexpression induced expression of Pparg but not Ppara, as well as PPARγ target genes such as *Cd36* and *Fabp4* as well as *Pdk4* which may participate in regulation of PDHA phosphorylation **(Figure 6C)**. Furthermore, HGFAC overexpression increased phosphorylation of PDHA (S293) as well as Proliferating Cell Nuclear Antigen (PCNA) levels, indicating increased proliferation **(Figure 6D)**. Whereas short-term overexpression of HGFAC was sufficient to produce glycemic and gene expression phenotypes reciprocal to *Hgfac* KO, we did not observe changes in hepatic or circulating triglyceride levels in this time frame (**Figure 6E**). Thus, HGFAC overexpression can induce changes in hepatic PPARγ expression and glucose homeostasis independently of its effects on hepatic lipids.

**Figure 6.**
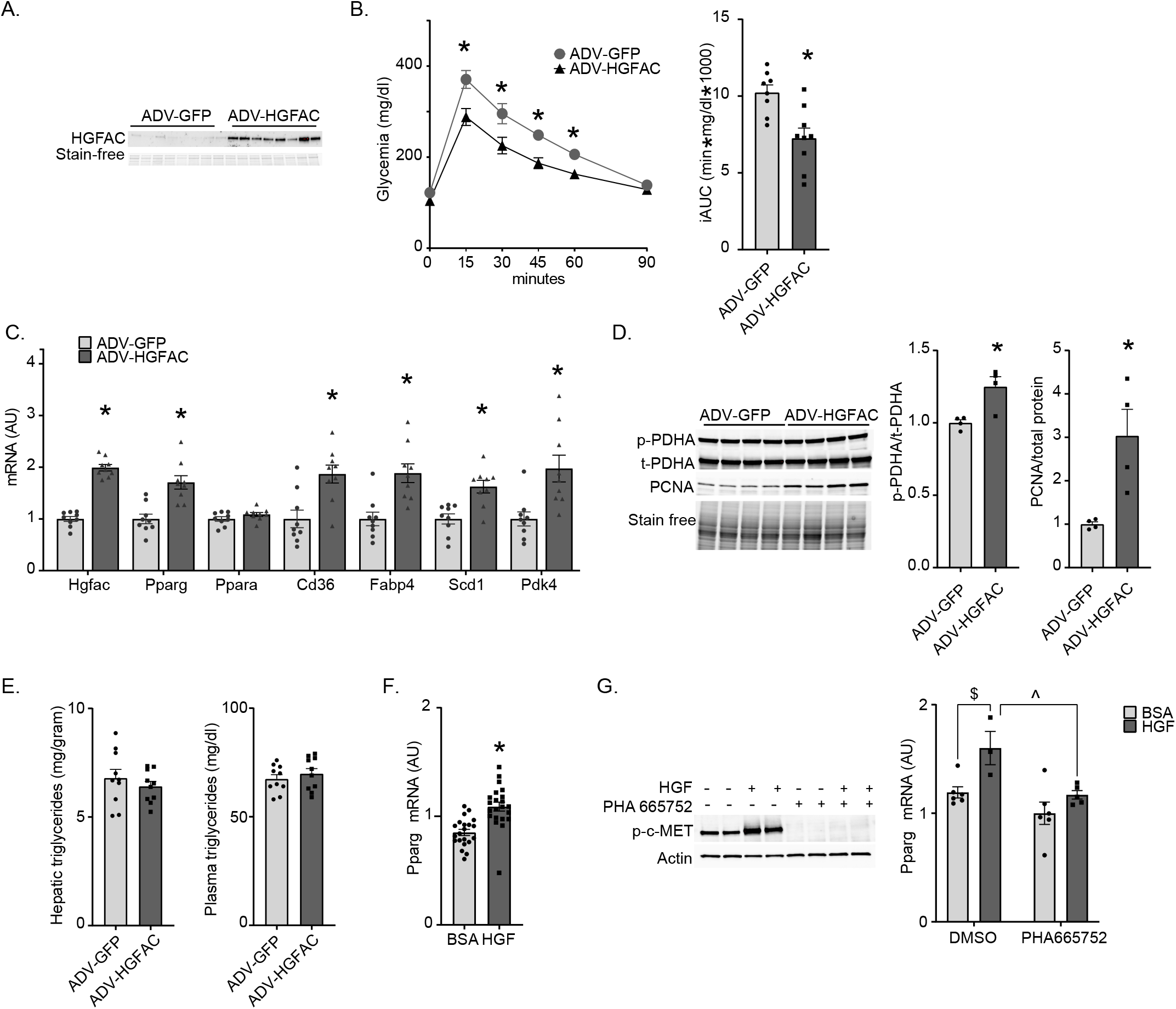
HGFAC overexpression enhances glucose homeostasis. A) Immunoblot of plasma HGFAC collected 3 days after 8-week-old male mice were transduced with adenovirus expressing GFP (ADV-GFP) or HGFAC (ADV-HGFAC). B) IP GTT and corresponding iAUC performed 5 days after viral transduction (n=10/group). C) hepatic mRNA levels of Hgfac, Pparg and -a and PPARγ targets measured by qPCR 10 days after viral transduction. D) Hepatic phospho-S293 PDHA, total PDHA, and PCNA immunoblots of liver from ADV-HGFAC and ADV-GFP transduced mice, and quantification of p-PDHA normalized to total PDHA and PCNA normalized to the total protein content, n=4 per group. E) Hepatic and circulating triglyceride levels 10 days after viral transduction in ad libitum fed mice. F) Pparg mRNA levels in AML12 cells after overnight treatment with 50 ng/ml HGF or BSA. G) c-MET phosphorylation by HGF in AML12 cells is inhibited by the c-MET inhibitor PHA 665752 preventing induction of *Pparg* mRNA. Data represent means ± SEM. Statistics assessed by two-tailed unpaired t-test, * p< 0.05; or by two-way ANOVA, ^ p<0.05 for comparison of effects of inhibitor within HGF treatment condition, $ p<0.05 for effect of HGF within inhibitor or control treatment.

To assess whether HGFAC’s effect to induce *Pparg* expression is likely mediated through its ability to activate HGF and c-MET signaling, we treated murine AML12 hepatocyte-like cells with recombinant, active HGF. HGF treatment increased c-MET phosphorylation and increased *Pparg* mRNA expression by 30% **(Figure 6F)**. These effects were inhibited by pre-treatment with PHA665752, a c-MET inhibitor **(Figure 6G)** (51). Altogether, these results support a model whereby overnutrition enhances ChREBP-dependent upregulation of HGFAC which activates an HGF-PPARγ signaling axis to preserve systemic glucose homeostasis.

## Discussion

ChREBP is a critical nutrient sensing transcription factor that is activated by cellular carbohydrate metabolites and mediates genomic and physiological responses to overnutrition in key metabolic tissues including the liver. The precise mechanisms by which carbohydrates active ChREBP remain controversial (reviewed in (1)). Putative mechanisms include carbohydrate mediated translocation of ChREBP protein from the cytosol to the nucleus, alterations in ChREBP post-translational modifications, and/or allosteric effects of specific carbohydrate metabolites on ChREBP itself to enhance transactivation. We previously demonstrated that fructose gavage acutely and robustly activates ChREBP-dependent gene expression in mouse liver following short-term fasting (4). Here, we performed ChIP-seq for ChREBP following fructose gavage after a 5 hour fast to map the induction of ChREBP binding in mouse liver chromatin. We identified ~ 4000 ChREBP binding sites in livers from two mouse strains that appears similar to previous efforts (10). To our surprise, chromatin-bound ChREBP was readily detectable in fasted animals and no significant increase in binding was observed following fructose gavage. These results suggest that while fructose can acutely activate ChREBP-dependent gene transcription, this transactivation is achieved by ChREBP that is already bound to chromatin and indicates that carbohydrate-stimulated nuclear translocation and accumulation of nuclear ChREBP is not essential for the ability of carbohydrates to enhance ChREBP’s transcriptional activity. These results favor models suggesting that either carbohydrate-mediated post-translational modification or allosteric activation are key mechanisms that might stimulate ChREBP’s transcriptional activity.

Variants in the human ChREBP locus associate with pleiotropic biological traits at genome-wide or near genome-wide significance with a particularly strong association with hypertriglyceridemia. The transcriptional targets that mediate ChREBP’s pleiotropic biological effects remain incompletely defined. By mapping ChREBP genomic binding sites in mouse liver and integrating this with human genetics data, we identified candidate transcriptional targets that might contribute to ChREBP-mediated regulation of circulating lipids. While genes and loci in proximity to ChREBP binding sites were enriched for variants that associated with hypertriglyceridemia, of the thousands of hepatic ChREBP binding sites, in this analysis, only ~ 2% of such sites contributed to the enrichment. We anticipate that a relatively small and partially overlapping subsets of ChREBP gene targets may contribute to its regulation of other metabolic traits.

Of the candidate genes identified here, a small minority comprised liver derived circulating factors or “hepatokines” that might regulate metabolism systemically. We elected to focus further attention on *HGFAC* as a putative ChREBP-regulated hepatokine and demonstrated that circulating HGFAC is indeed nutritionally regulated in a ChREBP-dependent manner. Moreover, we have shown that it participates in an adaptive metabolic response to obesogenic diets in part through its effects to stimulate hepatic *Pparg* expression and transcriptional activity (**Figure 7**).

**Figure 7.**
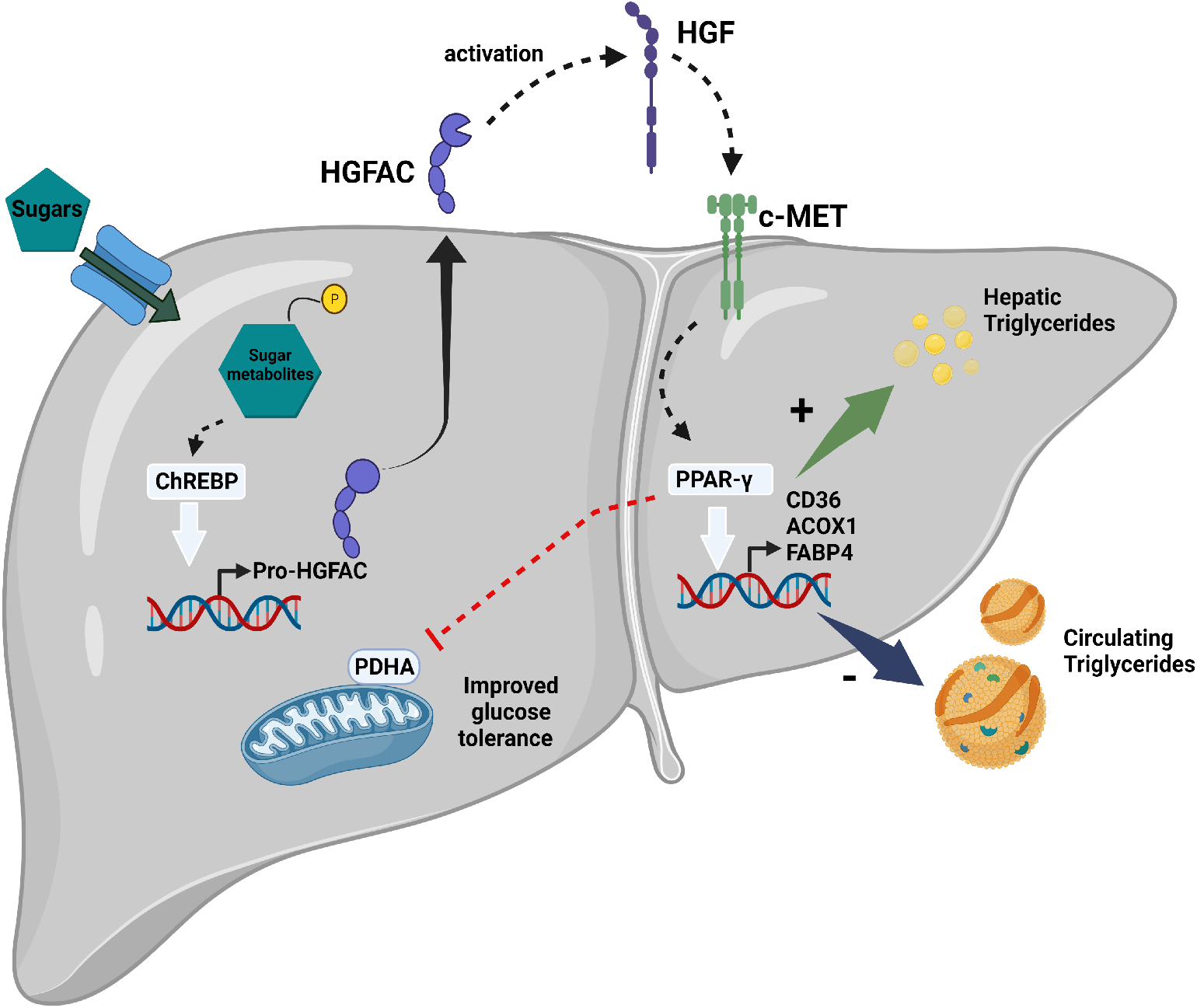
ChREBP mediated activation of a HGFAC-HGF-PPARγ signaling axis mediates an adaptive response to preserve glucose tolerance in the setting of diets high in sugar. Glucose and fructose from high sugar diets enhance production of sugar metabolites (hexose-phosphates) in the liver that activate hepatic ChREBP and lead to increased *Hgfac* transcription and translation. HGFAC is secreted into the circulation where, once activated, it can act in a paracrine or endocrine fashion to proteolytically cleave and activate HGF. HGF binds and activates the c-MET tyrosine kinase receptor on hepatocytes and other cell types. In liver, this leads to upregulation of PPARγ expression that in turn activates transcriptional programs to promote hepatic triglyceride storage and to decreased circulating triglycerides. Additionally, hepatic PPARγ activity decreases activation of the pyruvate dehydrogenase complex and this contributes to enhance systemic glucose tolerance.

To test the role of HGFAC in metabolism, we used a Crispr/Cas9 strategy to generate global *Hgfac* knockout mice. The ability of *Hgfac* KO serum to activate HGF and facilitate c-MET signaling was impaired. That this activity was attenuated, but not fully abrogated is consistent with known redundancy in enzymes capable of HGF activation (42, 52). Alternative proteases including kallikreins, urokinases, matriptase, and hepsin may partially compensate for loss of HGFAC activity (52–55). While the vast majority of HGFAC expression is in the liver, it is also expressed at much lower levels in other tissues including the intestines and possibly the pancreatic islets (23, 24). Therefore, we cannot exclude specific roles for HGFAC outside of the liver in this study.

The reduction in HGFAC activity and attenuation in HGF signaling did indeed produce metabolic phenotypes. We observed that *Hgfac* KO reduced and increased hepatic and circulating triglyceride, respectively and this was associated with impaired hepatic and systemic glucose tolerance. ADV-HGFAC overexpression produced a reciprocal phenotype with respect to glucose homeostasis but did not alter liver or circulating lipids in the short time frame of this experiment. Our results contrast with reported effects of acute treatment with recombinant, active HGF in rodents to reduce steatosis and with inconsistent effects on circulating triglycerides (56, 57). Additionally, marked and sustained transgenic overexpression of HGF under a metallothionein promoter also reduced steatosis in contrast with our observations (58). The differences observed in these publications and our experiments may be due to differences in gain-versus loss-of-function experiments, differential effects in acute versus chronic paradigms, and the degree of changes in HGF activity and signaling.

The specific mechanism by which pro-HGF is activated, either by HGFAC versus other proteases, also appears to have marked impact on where HGF signaling may be enhanced and on the resultant systemic metabolic effects. As an example, hepsin (HPN) is a membrane bound protease expressed in multiple tissues that is also capable of HGF activation. Hepsin KO which also reduces HGF-c-MET signaling produces a vastly different metabolic phenotype compared with *Hgfac* KO mice. Global hepsin KO mice are resistant to diet induced obesity and this lean phenotype is associated with enhanced glucose and lipid homeostasis (59). Profound changes in energy homeostasis in hepsin KO mice and its lean phenotype appear to be due to extensive expansion of brown fat and increased thermogenesis, features which we did not observe in *Hgfac* KO mice.

Proteases including HGFAC and HPN are promiscuous and may activate other peptide hormones which may also contribute to their differing biological effects. For instance, HGFAC can also cleave and activate pro-macrophage stimulating protein (pro-MSP, also known as MST1) which then activates the RON receptor tyrosine kinase (also known as MST1R) (60, 61). Although we determined HGF activation and c-MET signaling is impaired in experiments conducted with serum from *Hgfac* KO mice, it remains possible that some of HGFAC mediated changes that we observe are an effect of decreased signaling through MSP-RON cascade or other, unknown HGFAC proteolytic targets. Nevertheless, concordant associations in human HGFAC and MET variants with phenotype in *Hgfac* KO mice indicate that some of the key biological effects observed in *Hgfac* knockout mice are likely mediated through reduced HGF-MET signaling.

Our results show that the ChREBP-HGFAC axis regulates hepatic PPARγ signaling in mice. We further validated this observation by showing that HGF treatment can increase Pparg expression in hepatocyte-like AML12 cells and this can be blocked by a c-MET inhibitor. While the metabolic role of PPARγ is most well recognized with respect to adipogenesis, hepatic PPARγ also appears important in regulating systemic metabolism (62–64). Liver-specific deletion of Pparg reduces steatosis, but leads to hypertriglyceridemia and glucose intolerance associated with muscle and adipose insulin resistance (48). While the beneficial effects of hepatic PPARγ have been attributed to its effects on reducing circulating lipids, recent work has demonstrated that the PPARγ agonist pioglitazone enhances hepatic insulin sensitivity independently of its effects on hepatic lipids and is instead dependent on PPARγ’s ability to inhibit hepatic pyruvate dehydrogenase activity (50). Data from HGFAC KO mice are consistent with this hypothesis in that decreased PPARγ activity is accompanied by a reduction in inhibitory phosphorylation of the PDH catalytic subunit on Ser293. Adenoviral overexpression of HGFAC led to marked improvement in glucose tolerance with increased hepatic PPARγ expression and increased phosphorylation of PDH consistent with this model. Altogether, these results indicate that HGF and PPARγ may mediate some of its effects on glucose homeostasis through regulation of hepatic PDH phosphorylation.

Putative loss of function variants in human HGFAC strongly associate with increased circulating triglycerides, albumin, and platelets and these phenotypes are recapitulated in *Hgfac* KO mice (34). This concordance supports the hypothesis that putative HGFAC loss of function variants likely impair its catalytic activity. Moreover, these results suggest that this molecular physiology is conserved from rodents to humans. Interestingly, the rs1801282 (Pro12Ala) PPARG variant associated with increased *PPARG* expression and reduced risk for diabetes and circulating triglycerides also associates with reduced albumin levels (65). These effects on albumin are directionally concordant with the changes in albumin that occur in HGFAC KO mice and the reduction in hepatic *Pparg*. Again, this suggests that an HGF-PPARγ signaling axis is conserved in humans and that some of the beneficial effects of PPARγ on systemic metabolism could be mediated through effects in the liver in addition to adipose tissue.

Our results suggest an integrated physiology whereby carbohydrate sensing via ChREBP impacts systemic growth factor signaling (HGFAC-HGF-MET) that may mediate both adaptive and maladaptive responses through paracrine and endocrine effects. In the context of obesogenic diets, this signaling axis enhances hepatic *PPARG* expression which may mediate a compensatory response to preserve systemic glucose homeostasis. HGF, the principal target for HGFAC has previously been implicated in other aspects of glucose homeostasis. For example, HGF may enhance pancreatic beta cell proliferation (44, 66–68). Increased ChREBP-mediated HGFAC secretion might be a potential mechanism to increase beta cell mass in the setting of increased dietary carbohydrate burden. Additionally, within the liver, HGF has been reported to enhance insulin signaling and hepatic glucose clearance via physical interactions between its receptor, c-MET, and the insulin receptor (69). HGF also is secreted by adipocytes and can promote angiogenesis in adipose tissue and adipose angiogenesis is an integral feature of adipose tissue expansion (70–72). Therefore, elevated ChREBP-HGFAC-HGF may promote healthy expansion of adipose tissue for efficient storage of fuel during overnutrition. These observations may support a role for ChREBP mediated upregulation of HGFAC and HGF signaling as an adaptive response to increased nutritional burden, and will require further investigation. ChREBP itself has been shown to regulate mouse hepatocyte and murine and human beta cell proliferation (73–75). The ChREBP-HGFAC axis may provide an important mitogenic signal through HGF when ChREBP senses abundant carbohydrates indicative of ample building blocks supporting proliferation. Our data supports this hypothesis, as *Hgfac* KO animals have decreased expression of hepatic cell cycle genes, and adenoviral overexpression of HGFAC leads to marked upregulation of proliferating cell nuclear antigen in the liver, a marker of proliferation.

While a ChREBP-HGF signaling axis may mediate an adaptive response to overnutrition, both HGF and c-MET are well known oncogenes, and enhanced activation of HGF could conceivably potentiate oncogenesis (76, 77). Diets high in fructose, which activate hepatic ChREBP, are linked with increased incidence of hepatocellular carcinomas (HCC) in mice (78, 79). Furthermore, fructose and ChREBP have been implicated in development of non-alcoholic fatty liver disease both in animal models as well as in humans, which can progress to HCC in a subset of patients (80–83). Thus, ChREBP mediated signaling through HGF/c-MET might constitute an important link in the development of HCC in the setting of NAFLD/NASH.

Putative loss of function variants in HGFAC associate with increased circulating HGF in humans and also associate with increased cardiovascular risk factors (32, 34). Increased circulating HGF itself is increasingly recognized as a cardiometabolic risk factor that may be independent of other canonical cardiovascular risk factors (30, 32, 84–86). Further investigation into the relationship between ChREBP, HGFAC, and HGF signaling may define new mechanisms contributing to the pathogenesis of cardiometabolic disease in humans.

## Methods

### Reagents

Glucose (Cat #8769) and glycerol (Cat # G2025-1L) were purchased from Sigma, Ensure Original Nutritional Shake was purchased from a retail pharmacy, PHA-665752 (Cat # 14703) was purchased from Cayman Chemicals, mouse recombinant active HGF protein (2207-HG) was purchased from R&D systems, mouse Ultra-Sensitive Insulin Elisa was purchased from Crystal Chem Inc (Cat # 90080), Triglyceride LiquiColor test (Cat # 2200225) was from StanBio Laboratories. Thrombin (T4648-1KU) was from Sigma.

### Animals and diets

Floxed ChREBP mice were generated at UT Southwestern Medical Center as previously described (12). Albumin-Cre mice (stock 003574) were purchased from The Jackson Laboratory. Liver-specific ChREBP KO experiments were performed on a mixed C3H/HeJ and C57BL/6J background as previously described (3). *Hgfac* KO mice were generated at Duke Transgenic and Knockout Mouse Core, by introducing an 857 bp deletion from mid exon 1 and including exon 2 by CRISPR/CAS9 technology. All *Hgfac* KO experiments were performed on C57BL/6J background. Adenoviral overexpression of HGFAC was performed in wild type C57BL/6J male mice purchased from Jackson Laboratory (Bar Harbor). Mice were fed a chow diet (LabDiet 5008 or 5053), 60% fructose diet (TD.89247 Harlan Teklad) or 45% fat / 18% sucrose diet (D12451i, Research Diets) ad libitum for indicated times. All experimental mice were housed in 21°C–22°C on a 12-hour light-dark cycle in ventilated cages with 30 air exchanges per hour. Sequences for primers used for genotyping can be found in Supplemental **table 4.**

### Cell lines

AML12 (CRL-2254) and HepG2(HB-8065) cells were obtained from ATCC through the Duke Cell Culture Facility. AML12 cells were cultured in DMEM/F12 + 10% FBS supplemented with 1X Insulin-Transferrin-Selenium (ITS-G, 100X, Thermo Fisher, 41400045) and 40 ng/ml dexamethasone (Sigma, D4902). HepG2 cells were cultured in DMEM +10% FBS (Thermo Fisher, 16000044).

### ChIP-seq and Analysis

Wild-type, male 8-week old C3H/HeJ and C57BL/6J mice were fasted for 5 hours and gavaged with fructose (4 g /kg BW) versus water control (n=6 / group). Mice were euthanized 90 min after gavage and tissues were harvested and snap-frozen in liquid nitrogen for further analysis. Chromatin was prepared using truChIP Chromatin Shearing Tissue Kit (Covaris) according to the manufacture’s protocol with modifications. Briefly, 25-30 mg of frozen liver tissue were quickly minced with razor blades in PBS at room temperature. Tissue was crosslinked with 0.5 M disuccinimidyl glutarate in PBS for 45 min at room temperature, followed by second fixation with 1% formaldehyde in Fixing Buffer A (Covaris) for 5 min at room temperature. Crosslinking was stopped by Quenching Buffer E (Covaris). After washing, nuclei were isolated by Dounce homogenization followed by centrifugation. After washing and centrifugation, the nuclear pellet was resuspended in cold 0.25% SDS Shearing Buffer (Covaris). Chromatin was sheared in 1 ml AFA milliTUBEs (Covaris) using Covaris S220X focused ultrasonicator with the following parameters: peak incident power 140W, duty factor 5%, cycles per burst 200 for 12 min. The sheared chromatin was centrifuged at 13,000rpm for 10min at 4°C to remove debris, and a 10 ul aliquot was de-crosslinked and used for quantification with Qubit (Thermo Fisher Scientific). Sheared chromatin (1.5 – 3 μg) was diluted in ChIP dilution buffer (16.7 mM Tris [pH8], 1.2 mM EDTA, 25 mM NaCl, 1.1% Triton X-100, 0.01% SDS), and 1 μg of an ChREBP antibody (Novus, NB400-135) or control rabbit IgG was added, followed by overnight incubation at 4°C. The reactions were then incubated for 1h at 4°C with protein A/G dynabeads (Invitrogen) pre-blocked in PBS/0.5% BSA/0.5% Tween. Beads were then washed in low salt wash buffer (20 mM Tris [pH8], 1 mM EDTA, 140 mM NaCl, 1% Triton X-100, 0.1% sodium deoxycholate, 0.1% SDS), high salt wash buffer (20 mM Tris [pH8], 1 mM EDTA, 500 mM NaCl, 1% Triton X-100, 0.1% sodium deoxycholate, 0.1% SDS), LiCl wash buffer (10 mM Tris [pH8], 1 mM EDTA, 0.5% NP-40, 0.5% sodium deoxycholate, 250 mM LiCl) and TE buffer (10 mM Tris [pH8], 1 mM EDTA) and eluted and reverse cross-linked in elution buffer (10 mM Tris [pH8], 5 mM EDTA, 0.1% SDS, 300 mM NaCl, 0.8 mg/ml proteinase K, 10 μg/ml RNase A) by incubating at 65°C for 10 hours. Next, DNA was extracted using AMPure XP beads following the manufacturer’s manual and quantified by Qubit (Thermo Fisher Scientific). Immunoprecipitated chromatin was pooled by genotype and gavage condition for further analysis.

Library preparation, sequencing, and analysis were performed in the Boston Nutrition Obesity Research Functional Genomics and Bioinformatics Core. The sequencing ChIP-seq reads were demultiplexed using bcl2fastq and aligned to the GRCm38 mouse genome using Bowtie2 (87). PCR duplicates and low-quality reads were removed by Picard. Reads were processed using SAMtools and subjected to peak-calling with MACS2. SAMtools was also used to obtain 2 pseudoreplicates per sample (88, 89).

Only the peaks present in both pseudoreplicates were included for further downstream analysis. The coverage for peaks was obtained using BEDtools multicov (90). Normalization and differential analysis were performed using edgeR between fructose and water gavage conditions (91). To visualize ChIP-seq signals, reads were converted to the BigWig file format using BEDtools and bedGraphToBigWig (92). Peaks were tied to genes based on the nearest gene and transcription start site (TSS) within a radius of 200kb distance. The gtf file from GENCODE version M24 is filtered to include only processed transcript and protein coding transcript types as well as filtered for well supported transcripts (using only transcript support levels 1 and 2).

Additional analyses were performed as noted in the text. For MAGENTA analysis, genes included in the analysis were further filtered for transcriptional start sites which resided within 20 kb of a ChIP-seq peak. Human homologues of this filtered set of mouse genes were analyzed using the Meta-Analysis of Gene-set ENrichmenT of variant Associations (MAGENTA) algorithm in conjunction with joint Metabochip and GWAS triglyceride data from the global lipids genetics consortium (21, 27). Genes identified in this analysis were determined to be secretory proteins based upon their annotation in the UniProt database (93).

### Body weight and metabolic testing

Body weight was measured weekly. Circulating triglycerides were measured from ad libitum fed mice at 1pm, blood was collected from the tail vein. Body composition was measured by Bruker Minispec LF 90II. For glucose, glycerol, and insulin tolerance tests, mice were fasted for 5 hours starting at 7 am and glycerol or glucose were given via (2 g/kg body weight) via intraperitoneal injection. For Insulin tolerance tests, 1U insulin (Humulin R, Lilly) per kg was injected. Glucose measurements were performed using a handheld glucometer (Bayer Contour). For mixed meal tolerance test, mice were fasted overnight and gavaged with 10 ul/g of Ensure, blood was collected from tail vain at 0- and 10-minute time points for insulin measurement.

### Hepatic Triglyceride measurements

Liver neutral lipids were extracted with modified Folch method. Approximately 100 mg of liver tissue was homogenized in 3 ml Chloroform:methanol (2:1) and incubated with shaking overnight. After that 800 ul of 0.9% saline was added, vortexed, and centrifuged at 2000 G for 10 min. The chloroform phase was collected and dried overnight. Triglycerides were dissolved in butanol/Triton X-100/methanol (60/27/13 by volume) and measured using colorimetric triglyceride assay (StanBio).

### HGFAC/HGF activation assay

Blood was collected from 3 WT and 3 HGFAC KO mice and allowed to clot at room temperature for 1 hour, centrifuged at 7000 G for 15 min and serum was collected. Serum was incubated with 10 ug/ml dextran sulfate and 500 ng/ml of Thrombin 3 hour at 37°C. Activated serum was diluted with DMEM media (1:10). HepG2 cells were treated with this media for 5 min and then harvested.

Activation of c-MET was assessed by immunoblotting.

### Mouse Complete blood Count

Mouse complete blood count was performed with K2 EDTA treated plasma obtained from tail veins via an Element HT5 veterinary hematology analyzer (Duke University Veterinary Diagnostic Laboratory).

### Immunoblotting

Whole liver tissues were homogenized in lysis buffer containing 20 mM Tris-HCl, 150 mM NaCl, 1 mM Na_2_EDTA, 1 mM EGTA, 1% Triton, phosphatase (Pierce, A32957) and protease inhibitors (Sigma-Aldrich, P8340). Protein concentration was measured with BCA method (Thermo-Fisher, 23225). Approximately 40ug of protein was used for liver immunoblots. For plasma samples, 1ul of plasma was mixed directly with 15 ul of Laemmli buffer with reducing reagent added (NuPAGE™ Sample Reducing Agent, NP0004). Lysates were then subjected to immunoblotting with the indicated antibodies. Anti-HGFAC (R&D systems, AF1715), anti-β-Actin (Cell Signaling, 4970S), anti-phospho-c-MET (Cell Signaling 3077), anti-total c-MET (Cell Signaling 3127), anti-PDHA1 (phospho S293) antibody (Abcam, ab92696), Pyruvate Dehydrogenase (Cell Signaling 3205), anti-p85 (Upstate, 06-496) anti-PCNA (Cell Signaling, 2586). Quantification of blots were performed via imaging on a ChemiDoc XP (Bio-Rad) and use of associated Image Lab software v6.0. For loading normalization, whole lane proteins were quantified using Bio-Rad Stain free technology.

### qPCR

TRI reagent (Sigma, T9424) was used for RNA isolation from mouse liver and cell lines. RNA was reverse transcribed using a SuperScript VILO kit (Invitrogen). Gene expression was analyzed with the ABI Prism sequence detection system (SYBR Green; Applied Biosystems). Gene-specific primers were synthesized by Thermo-Fisher (**Supplementary table 4**). Each sample was run in duplicate, and normalized to *Tbp* (CHREBP LKO cohorts) or *Ppib* (HGFAC cohorts) RNA levels.

### Adenoviral overexpression of HGFAC in mice

Murine *Hgfac* cDNA was purchased from Sino biological (Cat: MG50039-M) and cloned via Gateway recombination into the pAd/PL-DEST (Thermo-Fisher, Cat # V49420) adenoviral vector with CMV promoter. The ADV-GFP control vector has been previously described in (94). Adenoviral vectors were produced and purified as previously described (94). Anesthetized mice were injected with 5*10^10 adenoviral particles expressing HGFAC versus GFP control via the retro-orbital route. Expression of HGFAC was assessed by immunoblotting plasma for circulating HGFAC three days after adenoviral transduction.

### RNA sequencing and analysis

RNA was isolated from mouse liver with TRI reagent (Sigma, T9424). RNA-seq was performed in Duke Center for Genomic and Computational Biology. RNA quality was assessed using a Fragment Analyzer (Advanced Analytical). mRNA capture, fragmentation, and cDNA library construction were conducted using a stranded mRNA-Seq Kit (Kapa biosystems, KR0960 – v6.17). 50bp paired-end sequencing was performed on an Illumina NovaSeq 6000 and at least 35 M reads were obtained per sample. Sequencing data were uploaded to https://usegalaxy.org/ and aligned with HISTAT2 (2.1.0) using Mus musculus (house mouse) genome assembly GRCm38 (mm10). Transcript levels were quantified using FeatureCounts(95). Transcript level count was uploaded to the BioJupies server and analyzed for differential gene expression and KEGG pathway enrichment (96–98).

### Human hepatic HGFAC gene expression and analysis

*HGFAC* mRNA expression values for lean, obese, and obese/diabetic patients were extracted from data deposited in Gene Expression Omnibus (accession no. GSE15653) (40). Liver RNA-seq read counts were obtained from the GTEX RNA-seq project (version 8, 2017-06-05). Genes with average expression value > 20 were selected, log transformed, and z-scores were computed. HGFAC correlation was calculated with every gene and sorted by Pearson correlation. The top 5% of correlated genes were selected and analyzed with enrichR against ARCHS4 transcription factor co-expression database (39, 99). For correlation between *HGFAC* and ChREBP targets, a composite expression vector for validated ChREBP targets (*PKLR, ALDOB, FASN, KHK* and *SLC2A2*) was computed by averaging the log transformed, z-score expression values for each of these genes.

### Statistics

All data are presented as the mean ± SEM. Data sets were analyzed for statistical significance with GraphPad Prism using two-tailed unpaired t-test, and where indicated 2-way ANOVA and with post-hoc comparisons performed with Sidak’s test or 1-way ANOVA with Holm-Sidak’s multiple comparisons test between control and individual groups. Statistical significance was set at P < 0.05.

### Study approval

All mouse studies were approved by the Beth Israel Deaconess Medical Center or the Duke University Medical Center Institutional Animal Care and Research Advisory Committee.

## Author contributions

MAH, AS, LD, SAH, and IA designed, performed, and interpreted mouse experiments. PJW performed rat experiments. MAH, LD, AS, WT, HS, RI and LT designed, performed, and interpreted computational analyses. AS and MAH designed, performed, and interpreted in-vitro experiments. JMH designed and performed construction of adenoviral vectors, and MA and PJW prepared purified adenoviruses, PAG, WT, RM and HHK assisted with performing and interpreting experiments. MAH conceived of, designed, and supervised the experimental plan, interpreted experiments. AS and MAH wrote the manuscript. All authors edited the manuscript.

## Acknowledgments

This work was supported by NIH grants R01DK100425 (MAH), 5R01DK121710 (MAH), P30DK057521 Pilot and Feasibility Award (MAH), American Heart Association 16CSA28590003 (MAH), P30DK04620-BNORC Functional Genomics and Bioinformatics Core (LT), American Diabetes Association 1–19-PDF-088 (AS), Pathway to Stop Diabetes Award from the American Diabetes Association 1-16-INI-17 (PJW), Borden Scholar Award (MAH), and Eli Lilly and Company (MAH). We thank the Duke University School of Medicine for the use of the Sequencing and Genomic Technologies Shared Resource, which provided RNA-seq service. Illustration created with BioRender.com.

