## Supplemental Figures for "HGFAC is a ChREBP Regulated Hepatokine that Enhances Glucose and Lipid Homeostasis"

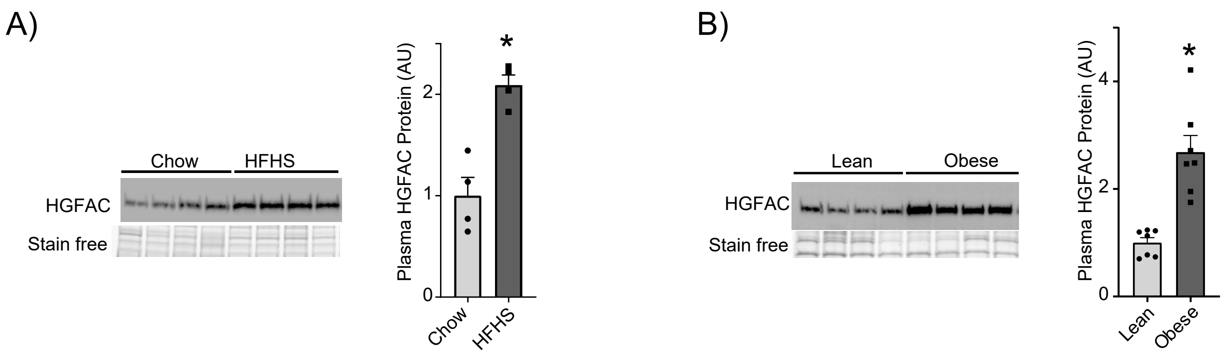


Supplementary figure 1. **Circulating HGFAC is increased in animals on obesogenic diets or with genetic forms of obesity.** Immunoblot analysis and quantification of circulating HGFAC in **(A)** male C57Bl/6J mice after 8 weeks on chow or HFHS diet, n=4 per group, and in **(B)** lean and obese Zucker rats, n=7 per group (Representative blot shown). Data represent means ± SEM, * $p<0.05$, two-tailed unpaired t-test.

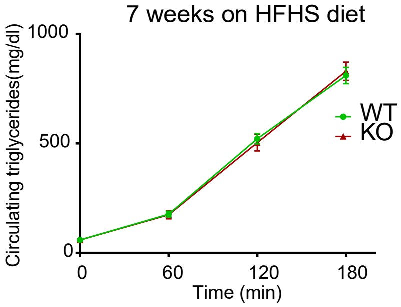


Supplementary Figure 3. **VLDL secretion is not different between HGFAC KO mice and controls (WT).** Plasma triglyceride levels in WT and Hgfac KO mice fed HFHS diet for 7 weeks after 3 hours food removal followed by injection with 1g/kg poloxamer 407 injection ip, n=12 per group.

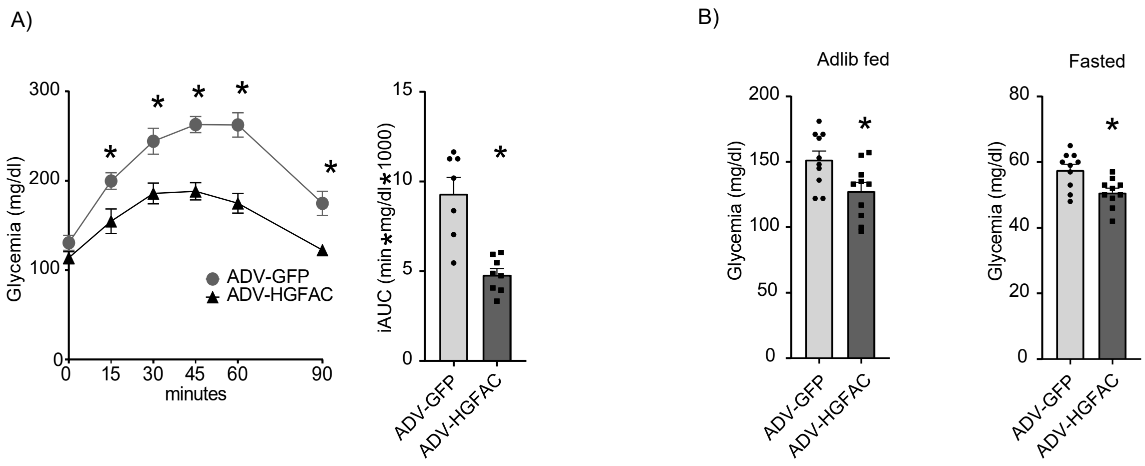


Supplementary Figure 5. **HGFAC overexpression leads to enhanced glucose homeostasis.** A) IP glycerol tolerance test and corresponding iAUC performed two weeks after viral transduction, (n=7-8/group). B) Ad libitum fed and 5 hour fasted peripheral glucose levels in ADV-GFP or ADV-HGFAC transduced mice. n=10 per group, Data represents Mean ± SEM, *p<0.05, two-tailed unpaired t-test.

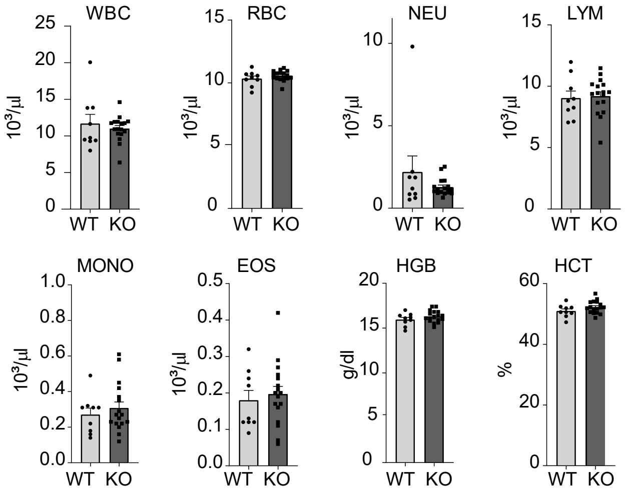


Supplementary Figure 2. **HGFAC KO does not influence blood parameters aside from platelet count.** Numbers of circulating white blood cells (WBC), red blood cells (RBC), neutrophils (NEU), lymphocytes (LYM), monocytes (MONO), and eosinophils (EOS), as well as hemoglobin (HGB) level and hematocrit (HCT) level are similar between Hgfac KO mice and controls (WT) n=9-17/g.

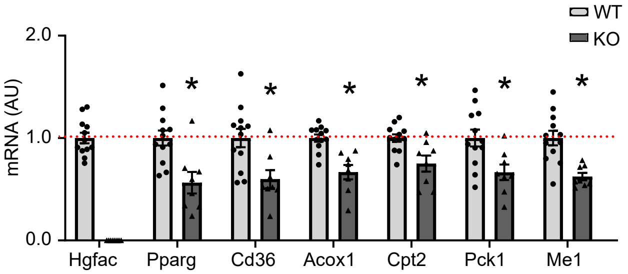


Supplementary Figure 4. **PPAR-γ and its targets are downregulated in HGFAC KO mice (replication cohort).** Hepatic mRNA levels of Hgfac, Pparg, Cd36, Acox1, Cpt2, Pck1 and Me1 in ad libitum chow fed Hgfac KO mice (KO) and controls (WT). n=8-12/group. Data represent means ± SEM, *p<0.05, two-tailed unpaired t-test.
